# Effects of rocuronium, sugammadex and rocuronium-sugammadex complex on coagulation in rats

**DOI:** 10.1101/656868

**Authors:** Ismar L. Cavalcanti, Alberto Schanaider, Louis Barrucand, Estêvão L. C. Braga, Nubia Verçosa, Hans D. de Boer, Luiz A. Vane

## Abstract

**Background:** Sugammadex is an alternative pharmacological drug that is capable of reversing neuromuscular blockades without the limitations that are presented by anticholinesterase drugs. Coagulation disorders that are related to treatment with sugammadex were reported. The exact mechanism of the effects on coagulation are not fully understood.

**Objective:** To evaluate the effects of rocuronium, sugammadex and the rocuronium-sugammadex complex on coagulation in an experimental model in rats.

**Design:** An experimental randomized animal study.

**Setting:** An experimental unit at the State University of São Paulo (UNESP), Botucatu, SP, Brazil.

**Interventions:** Wistar rats were randomly assigned into the following groups: the Control group; the Ssal Group – 0.5 mL of intravenous saline; the Sugammadex group – intravenous sugammadex (100 mg/kg); and the Rocuronium-Sugammadex group – intravenous solution with rocuronium (3.75 mg/kg) and sugammadex (100 mg/kg). Anaesthesia was performed by using isoflurane with controlled ventilation.

**Main outcome measures:** Coagulation factors were measured 10 minutes after the end of the preoperative preparation and 30 minutes after the administration of the drugs in accordance with the chosen groups.

**Results:** Platelet counts, prothrombin times and activated partial thromboplastin times were similar between the groups and between the moments within each group. There were reductions in the plasma fibrinogen levels between sample times 1 and 2 in the Rocuronium-Sugammadex group (*P* = 0,035).

**Conclusion:** The rocuronium-sugammadex complex promoted reductions in plasma fibrinogen counts, although the levels were still within normal limits.

## Introduction

A residual neuromuscular blockade is a practical problem in the field of clinical anaesthesia and may lead to adverse clinical outcomes, such as increases in pulmonary complications in the postoperative period, including pneumonia, atelectasis, hypoxemia and hypercapnia, which can result in increased morbidity and mortality [1, 2]. Neuromuscular monitoring and the routine use of reversal drugs are key factors in preventing postoperative residual neuromuscular blockades.

Sugammadex, which is a type of drug in a class of selective neuromuscular binding agents, is a modified γ-cyclodextrin that encapsulates the neuromuscular blocking drug (rocuronium or vecuronium) and results in the reversal of rocuronium- or vecuronium-induced neuromuscular block [3, 4]. Sugammadex was developed as an alternative pharmacological drug that would be capable of reversing neuromuscular blockades without the limitations that are presented by anticholinesterase drugs. It was first introduced by the scientific community in 2002; since then, both animal and human studies have demonstrated the efficacy and tolerability of this drug [5, 6]. Additionally, from the time that this drug was introduced, several adverse events that are related to the use of sugammadex, including coagulation disorders, have been reported [7–9]. *In vitro* experiments have demonstrated that sugammadex influences the effects of factor Xa in the coagulation pathway, which then leads to the inhibition of the prothrombine-to-thrombin conversion [10]. Furthermore, it may inhibit the production of factor Xa via the intrinsic pathway [10]. In human volunteers, it has been demonstrated that sugammadex at dosages of 4 mg/kg and 16 mg/kg resulted in moderate increases in prothrombin levels and activated partial thromboplastin times [10]. However, the exact mechanism for these effects on coagulation are not fully understood.

The aim of this study was to evaluate the effects of rocuronium, sugammadex and the rocuronium-sugammadex complex on coagulation in rats.

## Methods

The protocol and experimental procedures were approved by the Ethics Committee for Animal Experiments of the Botucatu Medical School - State University of São Paulo “Julio de Mesquita Filho” (UNESP).

Male Wistar rats (n = 28) weighing 300 - 500 g and provided by the Central Bioterium of Botucatu (UNESP) were included in the present study. The care of the animals in the study was performed according to the ARRIVE (animals in research: reporting *in vivo* experiments) statement.

### Experimental groups

In this study, the animals were randomly distributed into the following four groups (with seven animals in each group): the Control group that did not receive any anaesthetic intervention; the Ssal group received an administration of 0.5 mL IV saline without a drug infusion; the Sugammadex group received an intravenous administration of 100 mg/kg sugammadex (equivalent to a human dose of 16 mg/kg) [11] in 0.5 mL saline; and the Rocuronium-Sugammadex group received an intravenous administration of the combination of 3.75 mg/kg rocuronium (equivalent to a human dose of 0.6 mg/kg) [11] and 100 mg/kg sugammadex (equivalent to a human dose of 16 mg/kg) [12] in 0.5 mL saline.

In all of the experimental groups, the rats were anaesthetized in a closed container with a simultaneous flow of 2 L/min oxygen and 4% isoflurane during a time span of 5 - 10 minutes until the animals were unconscious. Thereafter, the trachea of each of the animals was intubated, and anaesthesia was maintained with 1.5% - 3% isoflurane in oxygen with a flow of 0.4 L/min. Controlled mechanical pulmonary ventilation was performed with a Fan Rodent Ventilator model 683™ (Harvard Apparatus, Holliston, United States). Inhaled anaesthesia and oxygen administration were performed by using the anaesthesia machine known as the Ohmeda Excel 210 SE™ (Datex-Ohmeda, Madison, United States), and the vaporizer was calibrated from the Isotec 5, Ohmeda™ (Datex-Ohmeda, Madison, United States).

### Data collection

The collection of haemodynamic parameters and blood samples was performed at two different time points (sample time): T1 was obtained at 10 minutes after the end of the preoperative preparation, and T2 was obtained at 30 minutes after randomly selected treatments with sugammadex, sugammadex-rocuronium or saline infusions in the respective groups. Liver and kidney tissue samples were immediately collected after the end of the experiment when all of the animals were euthanized with deep inhalation anaesthesia and intra-cardiac injections of 4 mL of 2.5% sodium thiopental.

### Experimental procedure

The following experimental sequences were conducted: 1 - solid food fasting for 2 hours with free access to water; 2 - weighing the animal; 3 - induction of anaesthesia by using 4% isoflurane in a closed container and spontaneous ventilation for 5 - 10 minutes; 4 - tracheal intubation; 5 - placement and fixation of the animals in a Claude Bernard device; 6 - mechanical ventilation with a tidal volume of 8 ml/kg of oxygen and isoflurane in concentrations that were sufficient to complete anaesthesia; 7 - dissection and cannulation of the left jugular vein for the purpose of hydration (5 mL/kg/h of saline solution that was administered by the use of an infusion pump [Anne™ manufacturer]); 8 - dissection and cannulation of the right carotid artery to monitor the heart rate (HR) and the mean arterial pressure (MAP); 9 - waiting 10 minutes after the preoperative preparation; 10 - measurements of haemodynamic and blood sampling parameters corresponding to time point T1. After the collection of arterial blood samples (2 mL), volume replacement was performed with 4 mL of intravenous saline; 11 - infusion of the appropriate drugs in the specific groups in accordance with the study protocol; 13 - 30 minutes after the infusion of appropriate drugs, measurements of the same parameters of T1 that corresponded to time point T2; after the collection of arterial blood samples (2 mL), volume replacement was performed with 4 mL of intravenous saline; 15 - animals euthanized; and 16 - end of the experiment.

### Blood sample and parameter collection procedures

HR and MAP data were obtained using a transducer known as the Datex angstrom model AS/3™ (Datex-Ohmeda, Madison, United States), which was attached to the carotid artery and allowed for the continuous monitoring of these data. The haematocrit data were obtained via blood sampling through the catheter, which was placed in the carotid artery, after which the collected blood was placed into two capillary tubes and transported to a microcentrifuge. The platelet counts were analysed by the use of fluorescence via an automatic flow cytometer (Beckman Coulter, São Paulo, Brazil). Prothrombin time (PT), activated partial thromboplastin time (APTT) and fibrinogen values were evaluated by the use of the photometric method via an automated coagulation analyser (MC-SMT100V, China).

### Statistical analysis

All of the samples were analysed using the Wilcoxon test, which showed that all of the data were parametric. The distributions of the data were normally distributed. For the independent variables, the Student’s t-test was used to compare the data.

For the fibrinogen levels that did not exhibit homogeneity of the variances, the Kruskal-Wallis test was performed when comparing the groups at both of the sample time points, and the Wilcoxon comparison was used for the comparison of the moments in each group. The level of significance was regarded as 0.05.

## Results

The animals in the Rocuronium-Sugammadex group exhibited higher weights than the animals in the other groups. However, the average weight remained within the predetermined inclusion parameters (Table 1). There was a statistically significant difference between the Control and Rocuronium-Sugammadex groups (p = 0.003).

**Table 1.** Body Weights (g).

| Groups | Mean | SD | 95% Confidence Interval |  |
| --- | --- | --- | --- | --- |
|  |  |  | Lower | Upper |
| **C** | 365.7 | 23.7 | 343.8 | 387.6 |
| **Ssal** | 370 | 36.51 | 336.2 | 403.8 |
| **Sug** | 355.7 | 39.52 | 319.2 | 392.3 |
| **Roc -**<br>**Sug** | 440 | 47.61 | 396 | 484 |

The body temperature, heart rate and median arterial pressure values were similar among all of the groups and at both sample time points within each group.

Haematocrit levels were similar among the groups. However, haematocrit levels exhibited statistically significant differences between the two sample time points in all of the groups, except for the Sug group (Table 2).

**Table 2.**
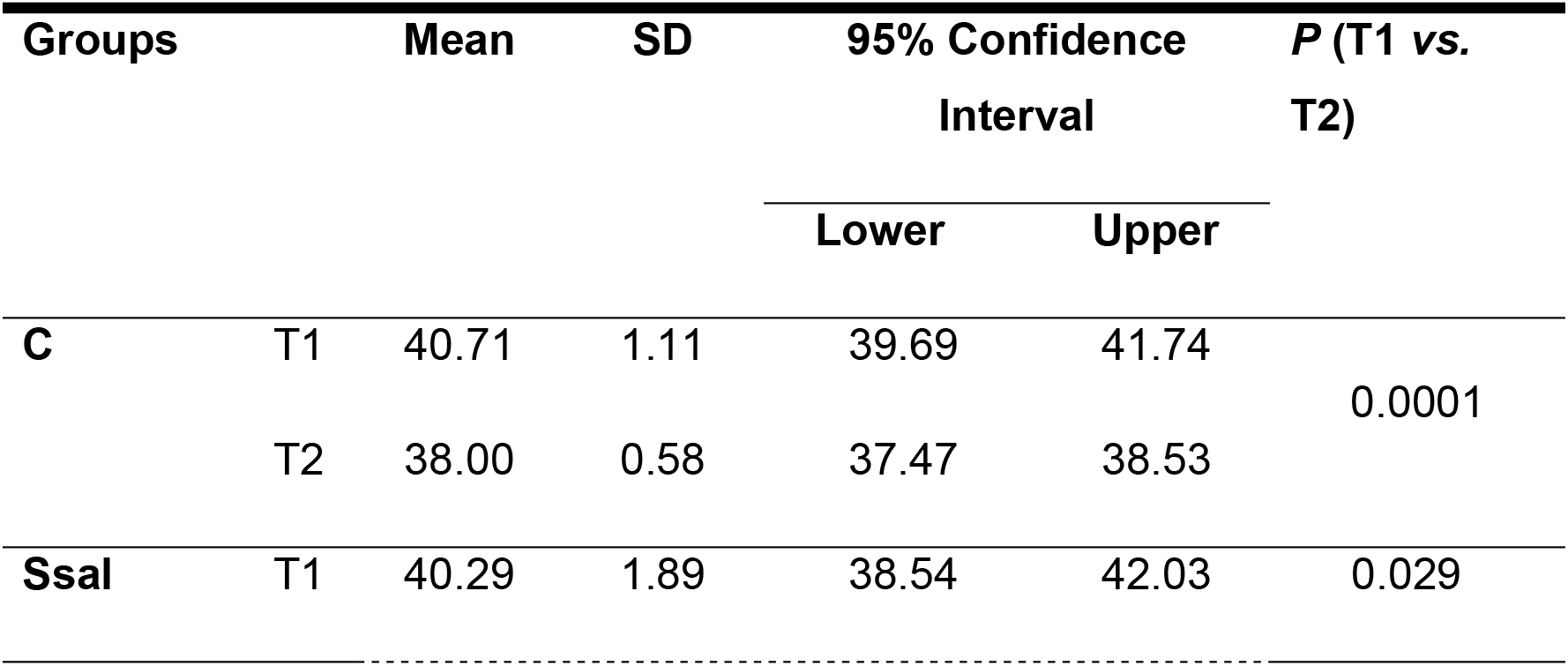
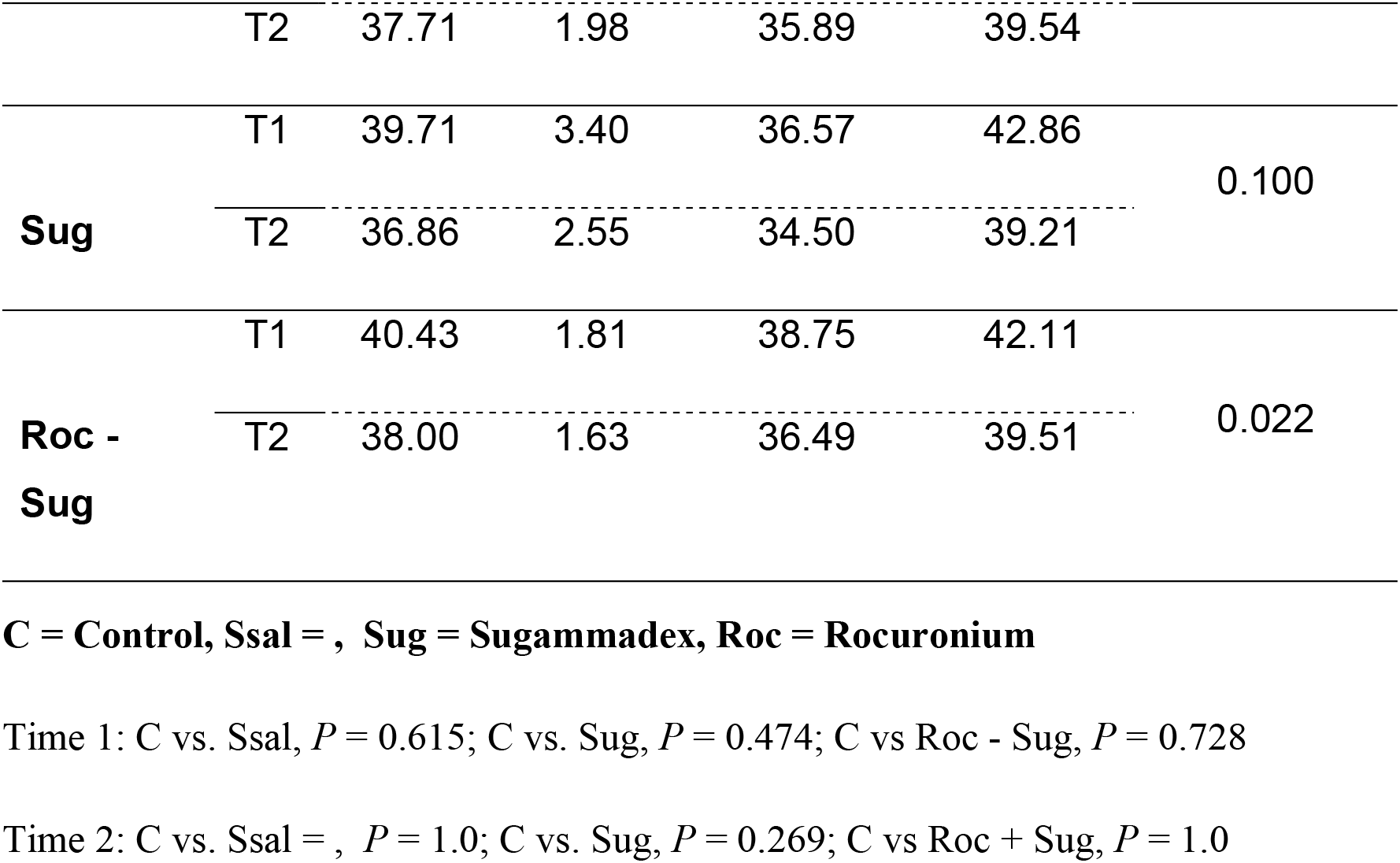
Haematocrit (%).

Platelet counts were similar among the groups and between the sampling time points within each group (Table 3).

**Table 3.**
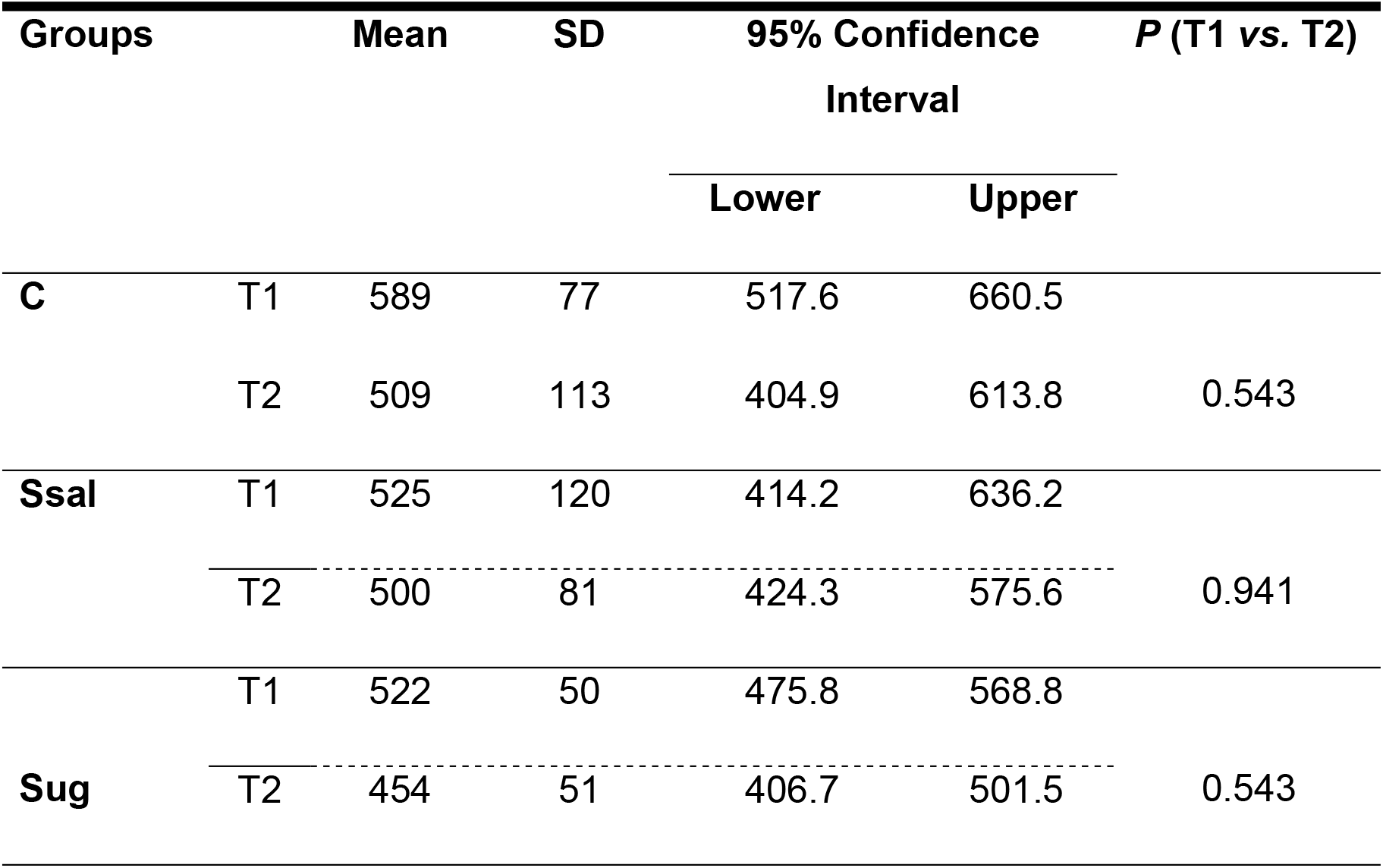
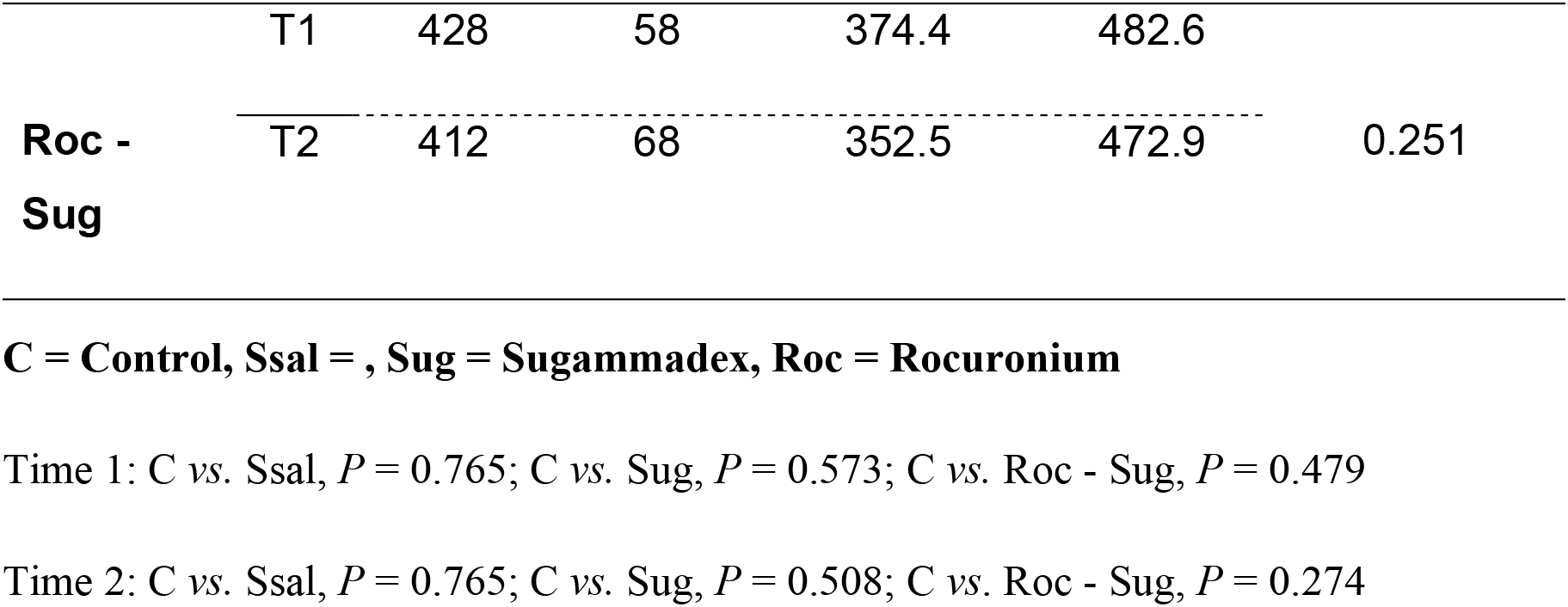
Platelet Counts (×10^3^/μL).

Prothrombin times were similar among the groups and between the sampling time points within each group (Table 4).

**Table 4.**
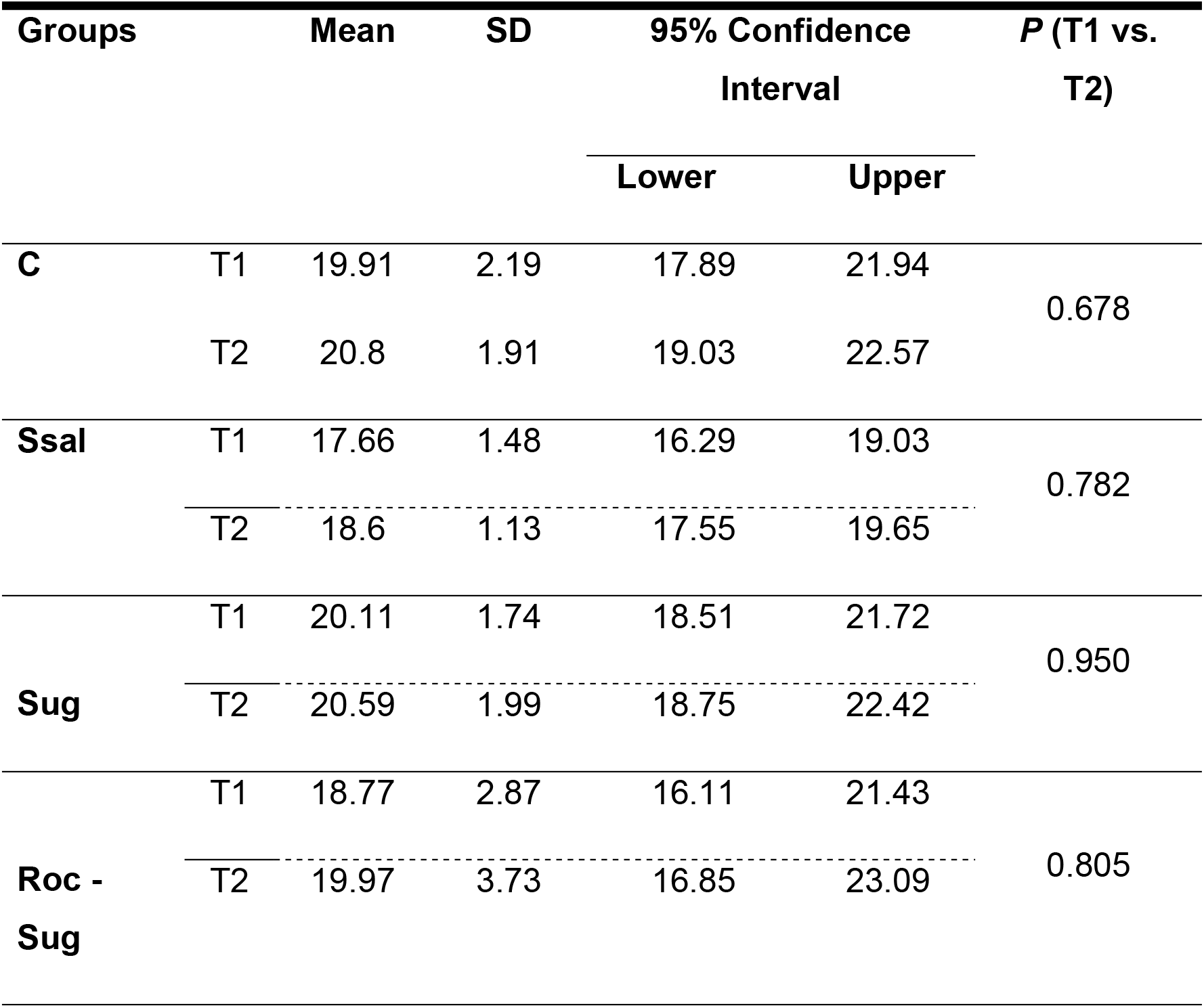
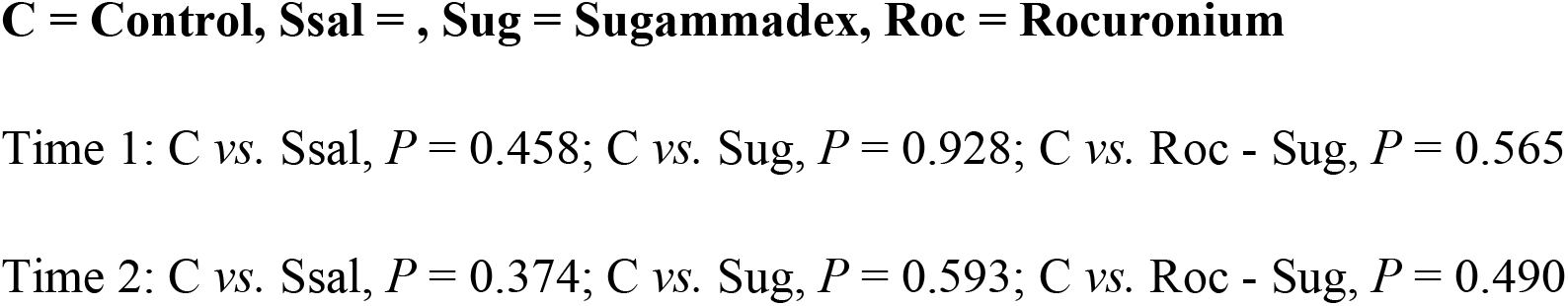
Prothrombin Times (s).

| Groups | | Mean | SD | 95% Confidence Interval | | $P$ (T1 vs. T2) |
| --- | --- | --- | --- | --- | --- | --- |
|  |  |  |  | Lower | Upper |  |
| **C** | T1 | 19.91 | 2.19 | 17.89 | 21.94 | 0.678 |
|  | T2 | 20.8 | 1.91 | 19.03 | 22.57 |  |
| **Ssal** | T1 | 17.66 | 1.48 | 16.29 | 19.03 | 0.782 |
|  | T2 | 18.6 | 1.13 | 17.55 | 19.65 |  |
| **Sug** | T1 | 20.11 | 1.74 | 18.51 | 21.72 | 0.950 |
|  | T2 | 20.59 | 1.99 | 18.75 | 22.42 |  |
| **Roc - Sug** | T1 | 18.77 | 2.87 | 16.11 | 21.43 | 0.805 |
|  | T2 | 19.97 | 3.73 | 16.85 | 23.09 |  |

Activated partial thromboplastin times (s) were similar among the groups and between the sampling time points within each group (Table 5).

**Table 5.**
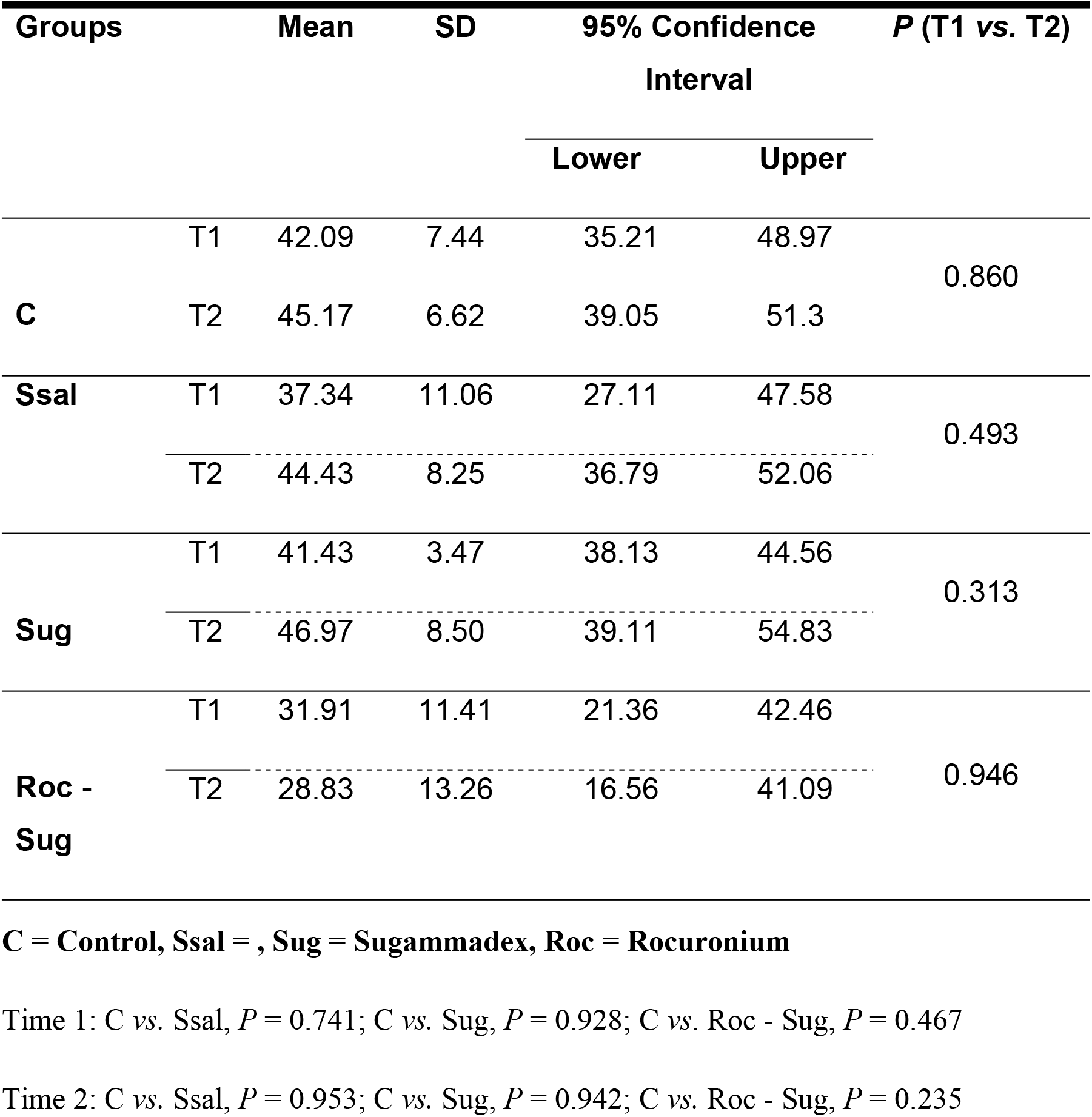
Activated Partial Thromboplastin Times (s).

There was a statistically significant reduction in plasma fibrinogen levels between sampling time points 1 and 2 in the Rocuronium-Sugammadex group (P = 0.035). There were no statistically significant differences among the other groups (Table 6).

**Table 6.** Plasma Fibrinogen (mg/dL).

| Groups | | Mean | SD | 95% Confidence Interval | | $P$ (T1 vs. T2) |
| --- | --- | --- | --- | --- | --- | --- |
|  |  |  |  | Lower | Upper |  |
| **C** | T1 | 250.7 | 59.89 | 195.3 | 306.1 | 0.618 |
|  | T2 | 285 | 166.7 | 130.8 | 439.2 |  |
| **Ssal** | T1 | 249.3 | 13.97 | 236.4 | 262.2 | 0.273 |
|  | T2 | 316.4 | 153.9 | 174.1 | 458.8 |  |
| **Sug** | T1 | 227.9 | 33.89 | 196.5 | 259.2 | 0.506 |
|  | T2 | 217.9 | 18.45 | 200.8 | 234.9 |  |
| **Roc - Sug** | T1 | 255 | 19.36 | 237.1 | 272.9 | 0.035 |
|  | T2 | 230 | 20 | 211.5 | 248.5 |  |

## Discussion

The main finding of this study was a reduction in the plasma fibrinogen levels in the Rocuronium-Sugammadex group.

In all of the groups that were examined, the weights of the animals were observed to be within the parameters of the inclusion criteria. None of the animals exhibited hypothermia, which could have influenced the pharmacodynamic effects of the neuromuscular blocking drugs [12].

The reduction in haematocrit levels at the T2 time point could be partially explained by the reduction of blood cells as a result of blood sample collection procedures, followed by the volume replacements with saline, thus resulting in a dilution of the blood cells [13].

The prothrombin time is used to evaluate the extrinsic pathway and the common pathway of coagulation (clotting factors II, V, VII, X and fibrinogen) [15]. Conceptually, the prothrombin time is increased in cases of deficiencies of fibrinogen or factors II, V, VII and X. Fibrinogen is a hexameric glycoprotein that is involved in the final stages of coagulation and the fibrin precursor manometers that are required for the formation of the platelet plug. Fibrinogen has a high molecular weight (number), is soluble in the plasma and is the active enzyme that mediates the thrombin-to-fibrin conversion [14].

The present study demonstrated a reduction in plasma fibrinogen in the Rocuronium-Sugammadex group. Although fibrinogen levels were reduced, they remained in the normal range, even without changes in the prothrombin time. The availability of fibrinogen is regulated through dynamic changes of synthesis and breakdown, in order to maintain coagulation function. Fibrinogen deficiency can lead to the dysfunctional polymerization of fibrin and can impair haemostasis, but the reduced levels of fibrinogen that were observed in this study did not lead to any bleeding in the preparations. However, haemodilution, hyperfibrinolysis, acidosis and hypothermia may result from the depletion of fibrinogen availability and may consequently impair the coagulation process [15].

Since no indications of acidosis or hypothermia and hyperfibrinolysis were present in the current study, it is assumed that the reduction of fibrinogen occurred as a result of haemodilution, due to the saline replacement of the blood volume that was collected for laboratory testing [13]. This may have led to a discrete haemodilution that may have promoted the reduction in fibrinogen levels. Therefore, the reduction in fibrinogen levels could be due to the methodological parameters of the study and not specifically because of the drugs that were examined.

The partial thromboplastin time corresponds to the time that it takes for the coagulation of recalcified plasma to occur in the presence of cephalin. The activated partial thromboplastin time is used to evaluate the efficiency of the intrinsic pathway of coagulation in the measurement of the formation of fibrin clots. A prolonged thromboplastin time indicates that one of the factors is presenting at a lower value than the normal value, or that there is the presence of inhibitors, such as factors VIII, IX, XI and XII (the intrinsic pathway) or factors II, V and X (the common pathway) [14]. The present study demonstrated that sugammadex did not change the activated partial thromboplastin time between the examined time points; therefore, after drug administration, the intrinsic coagulation pathway was preserved. Conversely, Rahe-Meyer et al. [16] demonstrated a transient (< 1 h) prolongation in the activated partial thromboplastin time after the administration of sugammadex. We can assume that this difference may be because our study was conducted in rats, whereas the research of Rahe-Meyer et al. [16] was performed in humans, and the time points of the measurements were different. Tag et al. [17] showed that the administration of sugammadex in rats shows no effects on PT, APTT and fibrinogen measurements, which were confirmed in this study.

A possible cause for the effect of sugammadex on the various coagulation assays has been previously described by Dirkmann et al. [18] and is explained by an apparent phospholipid-binding effect, thus suggesting that the anticoagulant effects of sugammadex are likely an *in vitro* artifact [18].

An important question to be asked is whether these adverse effects are reproducible in clinical settings of increased risks of bleeding in the postoperative period. Raft et al. [7] studied the clinical courses of postoperative coagulation disorders (reoperation for haemostasis and evaluation of bleeding in dressings) in human subjects and demonstrated that sugammadex was not associated with increased rates of bleeding. Meanwhile, Carron [19] claimed that definitive conclusions are not yet possible regarding bleeding risks in surgical patients receiving sugammadex. In a more recent *in vitro* study in healthy volunteers, it was suggested that supratherapeutic doses of sugammadex are associated with moderate hypocoagulation in whole blood, which could be confirmed via tromboelastography [20].

This study had some limitations. For example, due to the differences between species, it is not possible to extrapolate the results to humans. Moreover, in areas of clinical biochemistry, haematology and toxicology, there are considerable physiological variations between animal species, and even the positive results of this study may be due to methodological factors.

Further clinical studies are needed to not only investigate the overall influence of sugammadex on coagulation but also to examine the more specific influence of this drug on patients with coagulation disorders.

## Conclusions

In the present study on rats, the Rocuronium-Sugammadex complex group exhibited a reduction in plasma fibrinogen. However, the reduction in plasma fibrinogen levels remained within normal limits and did not lead to bleeding. Future clinical studies are needed in order to investigate the effects of sugammadex on coagulation, especially in patients with coagulation disorders.

